# *Caenorhabditis elegans* DAF-18/PTEN non-autonomously prevents tumors by enhancing calcium sensitivity

**DOI:** 10.64898/2026.08.31.748322

**Authors:** Jichao Deng, Ida Clémence Djiomo Mbieda, Armi M. Chaudhari, Olivier Gagné, Vincent Roy, Pier-Olivier Martel, Martin J. Simard, Patrick Narbonne

## Abstract

Insulin/IGF-1 signaling (IIS) centrally promotes stem/progenitor proliferation during development to translate nutrition into tissue expansion. In adults however, despite ongoing feeding and systemic IIS stimulation, most tissues stop growing. How adult tissues balance out IIS-induced growth is incompletely understood. Here, we report a direct molecular link between IIS and calcium responses that permits a global reduction of germ tissue turnover rates in spermless *C. elegans* hermaphrodites. We show that these spermless hermaphrodites require the key negative IIS regulator DAF-18/PTEN to prevent AKT-1,2/AKT from phospho-inhibiting the highly conserved small GTPase RHO-1/RHOA in their spermathecal necks to improve their calcium sensitivity. Their increased contractility restricts ovulation and triggers oocyte accumulation along with a concomitant downregulation of GSC proliferation, stabilizing their germline in a hyperplastic state. Similar IIS-calcium cross talks may explain how IIS promotes anabolism in adult tissues without causing their expansion, and why reduced PTEN activity provokes benign differentiated hamartoma-like tumors.

## Introduction

Insulin/IGF-1 signaling (IIS) centrally links nutrient uptake to stem cell growth/proliferation (Shim et al. 2013; Burchfield et al. 2025) and their differentiation (Lopez et al. 2013) into specialized cells, ensuring tissue expansion during development. In adults however, where tissue growth stops, additional mechanisms must exist to prevent IIS from inducing an excess of differentiated tissue in well-nourished individuals, apart from adipocytes. Despite limited examples, these homeostatic feedback loops rely on the secretion of various growth regulators by the stem cells’ differentiated progeny (Jiang et al. 2009; Mondal et al. 2011; Hsu et al. 2014) to relay information on tissue saturation and antagonize the anabolic effects of IIS (Narbonne et al. 2015; Valet and Narbonne 2022). Yet, these effects are not completely cancelled as IIS retains stem cell stimulatory power in most adult tissues (LaFever and Drummond-Barbosa 2005; Shim et al. 2013; Narbonne et al. 2015; Chidambaram et al. 2022; Cheng et al. 2024). If IIS keeps stimulating stem cell growth/proliferation in adult tissues, then how is their expansion prevented in well-fed adults?

The *C. elegans* germline offers a simple model to effectively address this question. In sperm-bearing adult wild-type hermaphrodites, germline stem cell (GSC) proliferation is stimulated by both IIS and MPK-1/ERK activities to sustain rapid oocyte production (Lee et al. 2007; Narbonne et al. 2015; Narbonne et al. 2017; Robinson-Thiewes et al. 2021). As sperm is required to release oocytes from diakinesis and induce ovulation, arrested oocytes accumulate in the proximal gonad of spermless or unmated feminized hermaphrodites (Miller et al. 2001), until a homeostatic feedback loop inhibits GSC proliferation/differentiation to pause oocyte production (Morgan et al. 2010; Narbonne et al. 2015; Cinquin et al. 2016). This signal downregulates MPK-1/ERK activity to oppose IIS actions and downregulate GSC proliferation, altogether slowing down the production of new oocytes to match their now much slower ovulation rates (McCarter et al. 1999). Thus, a drop in oocyte needs stabilizes the germline in a hyperplastic state, where several oocytes can accumulate in an organized manner to give these spermless hermaphrodites a head start on reproduction before mating.

We previously showed that DAF-18, the *C. elegans* ortholog of the Phosphatase and TENsin homolog PTEN, a dual-specificity phosphatase best known for antagonizing IIS activity, was essential to downregulate oocyte turnover in spermless hermaphrodites. Namely, oocytes spontaneously activated and were ovulated in feminized *fog-1; daf-18(ø)* double mutants, while their GSCs kept proliferating to sustain, in vain, a full-blown oogenic program (Narbonne et al. 2015; Narbonne et al. 2017). Here, we show that DAF-18/PTEN surprisingly accomplishes those feats entirely cell non-autonomously, by promoting contractility in this organism’s spermathecal (Sp) neck myoepithelium to prevent the spontaneous entry of oocytes, ensuring their meiotic arrest and accumulation. This permits a homeostatic signal to prevent unnecessary GSC proliferation, altogether slowing down germ tissue turnover in response to sperm unavailability. Mechanistically, we find that DAF-18/PTEN promotes Sp neck contractility by increasing internal calcium (Ca^2+^) sensitivity, something it accomplishes through preventing AKT-1,2/AKT from directly phospho-inhibiting the small GTPase RHOA ortholog RHO-1 on a highly conserved site. Altogether, our results show that DAF-18/PTEN can link stem cell proliferation rates with differentiated cell needs entirely cell non-autonomously, while we identify a novel and potentially conserved key IIS-Ca^2+^ molecular crosstalk that could have much wider implications.

## Results

### DAF-18/PTEN non-autonomously slows down germ tissue turnover from the soma

In control (spearm-bearing) adult hermaphrodites, DAF-18::mNG is ubiquitous and present at oocyte membranes (Figure 1A) (Masse et al. 2005; Suzuki and Han 2006; Brisbin et al. 2009; Liontis et al. 2026). In hyperplastic feminized (spermless) *fog-1* mutants however, markedly higher DAF-18::mNG levels were present at the membrane of arrested oocytes (Figure 1A). Given this expression pattern and DAF-18’s importance for preventing MPK-1/ERK activation in the proximal oocytes of spermless hermaphrodites (Suzuki and Han 2006; Brisbin et al. 2009), we hypothesized that germline DAF-18 autonomously induced oocyte quiescence/accumulation in the absence of sperm. We therefore introduced a germline-specific *daf-18*-rescuing single-copy transgene (Frokjaer-Jensen et al. 2008; Dickinson et al. 2013), henceforth referred to as *germline::DAF-18*, in the genome of *fog-1; daf-18(ø)* mutants. Since *daf-18(ø)* mutant larvae are maternally rescued for dauer entry (Gil et al. 1999), the larval progeny of *germline::DAF-18* animals may carry diluted somatic DAF-18 activity in addition to robust germline activity (Figure 1B). Unexpectedly, the resulting transgenics continued to waste their oocytes like their *fog-1; daf-18(ø)* parents. Moreover, GSC proliferation (as inferred from the GSC mitotic index [MI]) (Crittenden et al. 2006; Narbonne et al. 2015; Narbonne et al. 2017; Robinson-Thiewes et al. 2021) stayed high in *fog-1; daf-18(ø); germline::DAF-18* adults, as in wild-type and *fog-1; daf-18(ø)* controls (Figures 1C-1F). To validate that our *germline::DAF-18* rescue transgene was functional and confirm these surprising negative results, we asked if it rescued dauer development in *daf-2; daf-18(ø)* dauer-defective double mutants, since maternally provided DAF-18 is sufficient to promote dauer entry (Gil et al. 1999). As expected, *germline::DAF-18* maternally rescued dauer formation in a *daf-2; daf-18(ø)* background (Figure 1G). These results suggest that germline DAF-18 cannot compensate for the ovulation and GSC proliferation defects of *daf-18(ø)* mutants.

**Figure 1.**
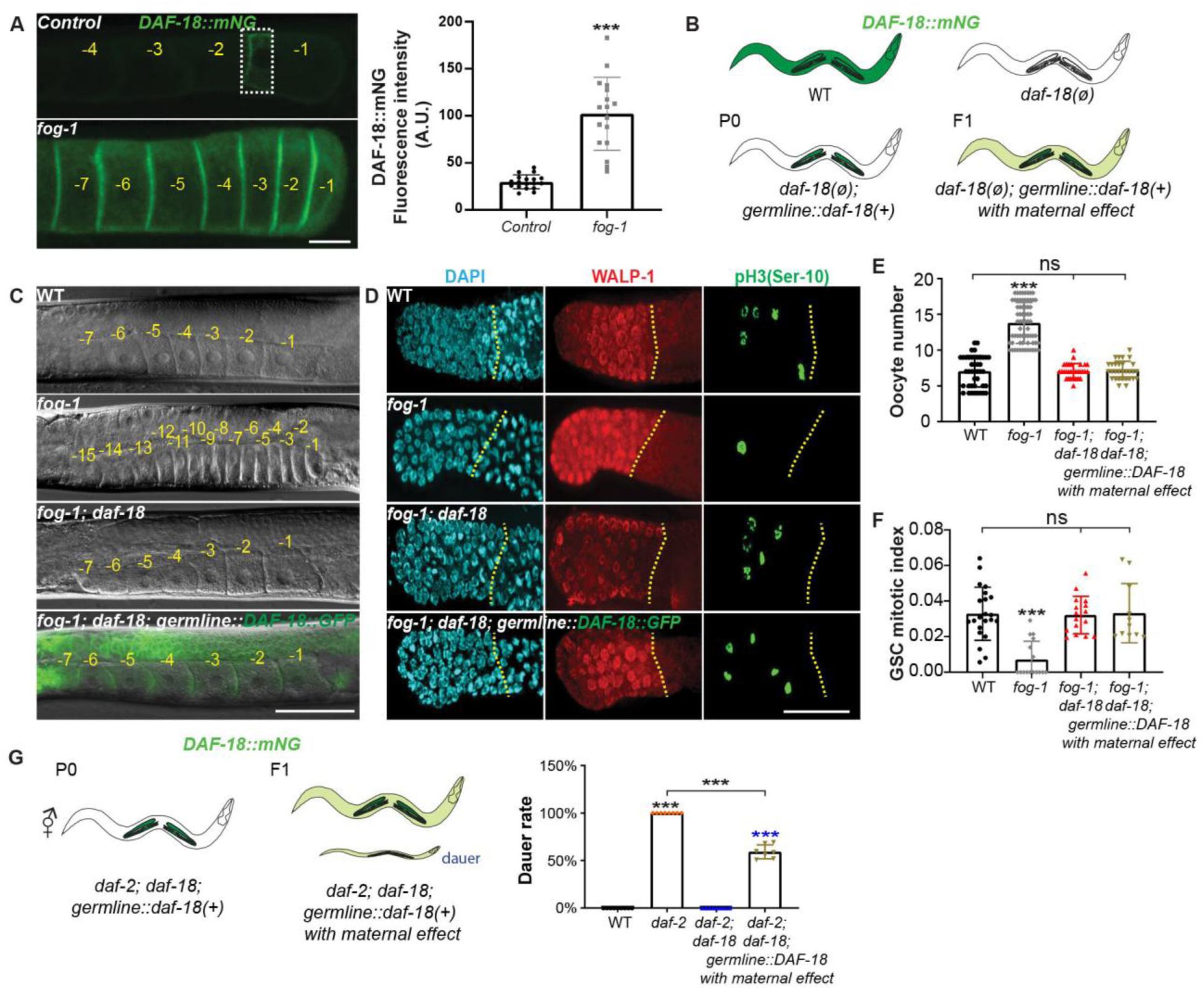
DAF-18/PTEN slows down germ tissue turnover cell non-autonomously from the soma. **(A)** Representative epifluorescence micrographs (left) and quantification (right) showing increased levels of endogenous tagged *DAF-18::mNG* at oocyte membranes in *fog-1* A1 hermaphrodites (see methods). Micrographs were acquired using identical parameters, but signal in the boxed region was enhanced in the control for visualization purposes. Sample sizes: 18, 15. **(B)** Schematic representation of the *germline::DAF-18* transgenic rescue experiment, showing expected pattern and approximate intensities of DAF-18::mNG in shades of green. **(C)** Representative differential interference contrast (DIC) or DIC/epifluorescence micrographs of A1 hermaphrodites of the indicated genotypes. **(A, C)** Negative yellow numbers mark oocytes from distal to proximal. **(D)** Representative epifluorescence micrographs of distal gonads dissected from A1 hermaphrodites of the indicated genotypes. Gonads were stained with DAPI (DNA; blue), anti-WALP-1 (proliferation maker; red) and anti-phospho[ser10] histone H3 (G_2_/M-phase marker; green). Yellow dotted lines, proliferative zone (PZ) boundary. Distal, left. (**A, C, D**) Scale bars: 50 µm. **(E)** Average number (± standard deviation) of diakinesis-stage oocytes per gonad arm in A1 hermaphrodites of the indicated genotypes. Sample sizes: 42, 68, 30, 29. **(F)** Average GSC MIs of A1 hermaphrodites of the indicated genotypes. Sample sizes: 22, 16, 16, 11. **(G)** Germ-specific DAF-18(+) expression rescues dauer formation in *daf-2; daf-18(ø)* mutants. Sample sizes: 9, 8, 9, 7. **(A,E-G)** Black asterisks indicate statistical significance from all other samples or as indicated by brackets, and colored one versus the corresponding color-coded sample. ns, not significant.

Further experimenting with this *germline::DAF-18* transgene revealed that zygotic germline expression was insufficient to promote dauer entry in a *daf-2; daf-18(ø)* background (Figure S1A), and that it maternally rescued dauer formation in an allelic dosage-dependent manner (Figure S1B). However, *germline::DAF-18* did not prevent the GSC overproliferation that occurs during dauer development in *daf-18(ø)* mutants (Figure S1B), something that requires somatic gonad DAF-18 activity (Tenen and Greenwald 2019). Altogether, these results are consistent with germ-specific transgenic expression and establish that the oocyte quiescence defects of feminized *daf-18(ø)* mutant adults are strictly zygotic. Since the maternal contribution from *germline::DAF-18* to all somatic tissues was insufficient to ensure oocyte arrest and inhibit GSC proliferation (Figures 1C-1F), we conclude that slowing germ turnover in the absence of sperm requires significant amounts of DAF-18 protein within somatic tissues.

### Muscle DAF-18 is sufficient to slow down germ tissue turnover

To identify the somatic tissue(s) from which *daf-18* slows down germ turnover (suppresses both ovulation and GSC proliferation) in the absence of sperm, we restored DAF-18 activity in each of the main somatic compartments of *fog-1; daf-18(ø)* doubles. We used the *unc-54, nhr-72, elt-7,* and *unc-119* promoters to specifically drive *daf-18* expression in non-pharyngeal muscles, seam cells, intestine and nervous system, respectively (Okkema et al. 1993; Maduro and Pilgrim 1995; Miyabayashi et al. 1999; Masse et al. 2005). Expression of DAF-18 in muscles restored oocyte arrest/accumulation and suppressed GSC proliferation, while expression in hypodermal seam, intestinal or neuronal cells did not; results that were confirmed with a second muscle-specific promoter (*Pmyo-3*) (Okkema et al. 1993) (Figures 2A-2D). In the absence of sperm, the presence of DAF-18 in non-pharyngeal muscles therefore non-autonomously ensures oocyte arrest, leading to their hyperplastic accumulation and the concomitant downregulation of GSC proliferation.

**Figure 2.**
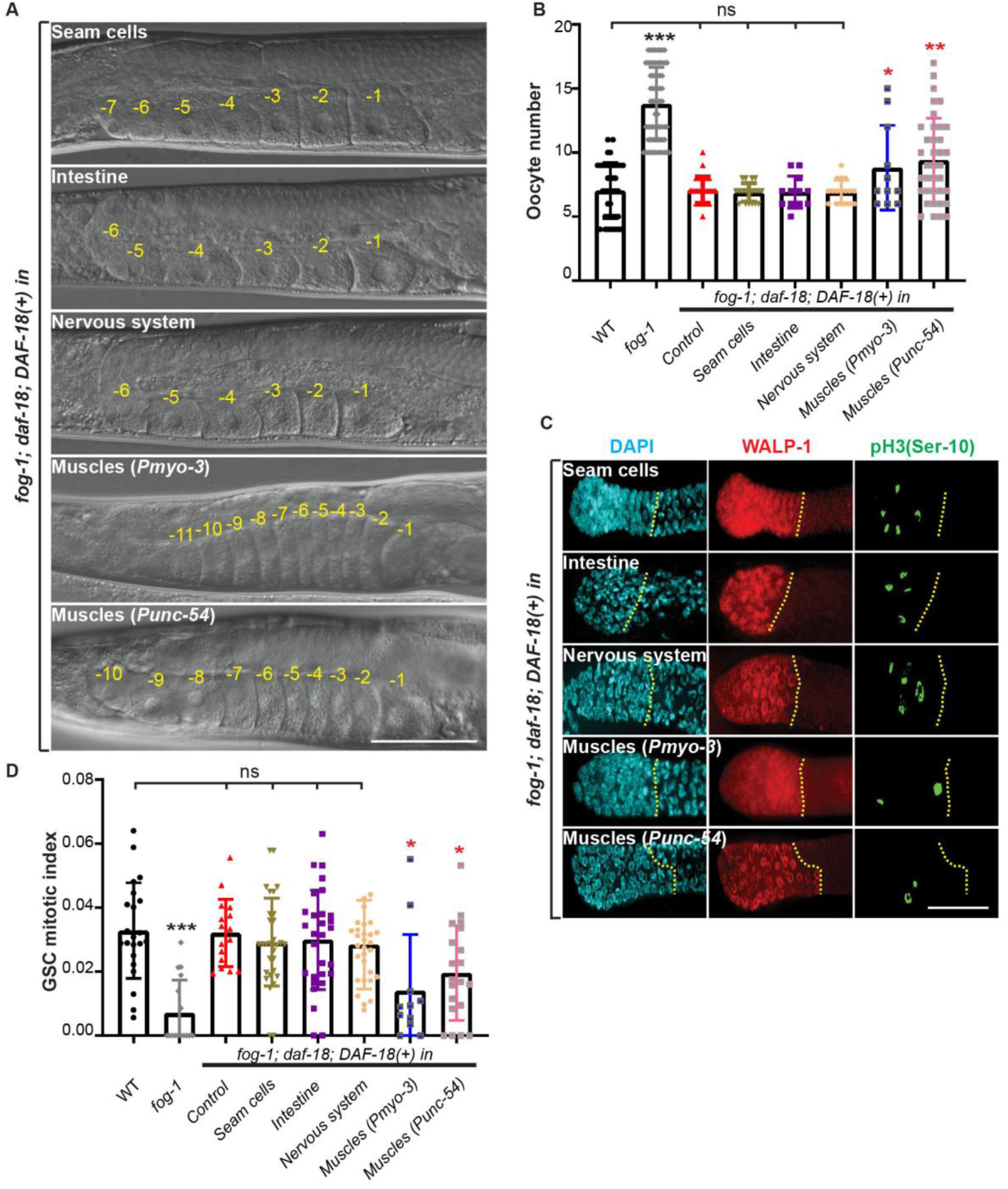
Muscle DAF-18 is sufficient to slow down germ tissue turnover. **(A)** Representative DIC micrographs of A1 hermaphrodites of the indicated genotypes. Negative yellow numbers mark oocytes from distal to proximal. **(B)** Average number (± standard deviation) of diakinesis-stage oocytes per gonad arm in A1 hermaphrodites of the indicated genotypes. Sample sizes: 42, 68, 30, 19, 12, 17, 11, 37. **(C)** Representative epifluorescence micrographs of distal gonads isolated from A1 hermaphrodites of the indicated genotypes, stained with DAPI (DNA; blue), anti-WALP-1 (proliferation maker; red) and anti-phospho[ser10] histone H3 (G_2_/M-phase marker; green). Yellow dotted lines, proliferative zone (PZ) boundary. Distal, left. (**A, C**) Scale bars: 50 µm. **(D)** Average GSC MIs of A1 hermaphrodites of the indicated genotypes. Sample sizes: 22, 16, 16, 31, 29, 30, 11, 20. **(B, D)** Black asterisks indicate statistical significance from all other samples and colored ones versus the corresponding color-coded sample. ns, not significant.

### Spermathecal neck DAF-18 is sufficient to slow down germ tissue turnover

The hermaphrodite muscular system consists of 20 pharyngeal and 95 body wall muscle cells, in addition to a few other specialized muscles (Altun and Hall 2009). While both the *myo-3* and *unc-54* promoters are not active in pharyngeal muscles (Miller et al. 1986; Ardizzi and Epstein 1987), it was conceptually difficult to hypothesize how body wall muscles could promote oocyte arrest in the absence of sperm. In contrast, the Sp and gonadal sheath cells are smooth muscle-like contractile cells that have been heavily implicated in the control of oocyte maturation and ovulation (McCarter et al. 1999; Miller et al. 2001). We closely examined animals bearing *myo-3* and *unc-54* transgenes and detected expressions from both promoters in the uterus (Ut), Sp and gonadal sheath cells (Figure S2). To determine whether *daf-18* may act within these contractile gonadal tissues, we first restored it specifically in the sheath cells, or in the proximal gonad comprising the Sp and Ut, of *fog-1; daf-18(ø)* doubles using the *lim-7* and *fos-1a* promoters, respectively (Figure 3A) (Voutev et al. 2009; Qin and Hubbard 2015). Only *Sp+Ut::DAF-18* expression restored oocyte accumulation and reduced GSC proliferation (Figures 3B-3E). Interestingly, while *sheath::DAF-18* had no effect on oocyte accumulation, it partially rescued GSC downregulation (Figures 3B-3E). This mild decrease in GSC proliferation in the absence of oocyte accumulation suggests that sheath cells are not the primary site for DAF-18 function. Moreover, as sheath MPK-1A activity can promote GSC proliferation, while DAF-18 can suppress MPK-1 activation (Suzuki and Han 2006; Nakdimon et al. 2012; Narbonne et al. 2017), this partial rescue likely results from the inhibition of sheath MPK-1A by DAF-18 overexpression. We infer that the Sp and Ut comprise the main site where DAF-18 activity downregulates both ovulation and GSC proliferation in spermless hermaphrodites.

**Figure 3.**
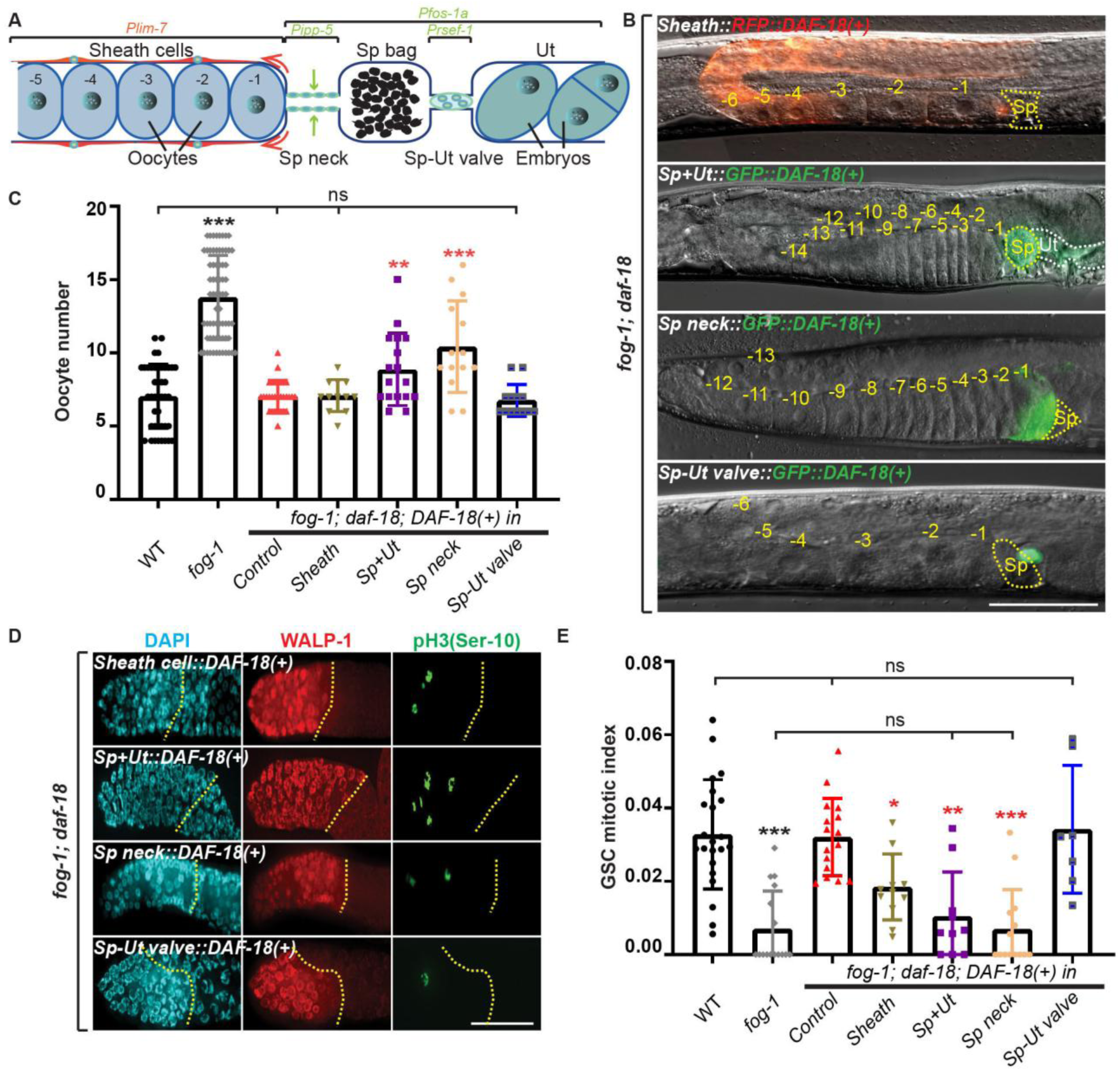
DAF-18 slows down germ tissue turnover from the spermathecal neck. **(A)** Schematic representation of the *C. elegans* proximal germline and somatic gonad. Regions of expression from tissue-specific promoters are indicated by brackets at the top. **(B)** Representative DIC/epifluorescence micrographs of A1 hermaphrodites of the indicated genotypes. Negative yellow numbers mark oocytes from distal to proximal. Yellow dotted lines, Sp. White dotted line, Ut. Dorsal, up; proximal, right ventral. **(C)** Average number (±standard deviation) of diakinesis-stage oocytes per gonad arm in A1 hermaphrodites of the indicated genotypes. Sample sizes: 42, 68, 30, 12, 13, 17, 14. **(D)** Representative epifluorescence micrographs of distal gonads isolated from A1 hermaphrodites of the indicated genotypes, stained with DAPI (DNA; blue), anti-WALP-1 (proliferation maker; red) and anti-phospho[ser10] histone H3 (G_2_/M-phase marker; green). Distal, left. Yellow dotted lines, PZ boundary. (**B, D**) Scale bars: 50 µm. **(E)** Average GSC MIs of A1 hermaphrodites of the indicated genotypes. Sample sizes: 22, 16, 16, 12, 7, 10, 14. **(C, E)** Asterisks indicate statistical significance versus the corresponding color-coded sample. ns, not significant.

The Sp is the site of fertilization, and consists of three parts: an 8-cell distal neck, a 16-cell central bag, and a syncytial 4-cell Sp-Ut valve (Figure 3A) (McCarter et al. 1997). To materialize ovulation, the gonadal sheath cells enter in a tug-of-war with the Sp neck until they successfully pull it open around the proximal (-1) oocyte (Castaneda et al. 2020). Once the oocyte enters the Sp, it is immediately fertilized, and the Sp neck and bag then contract to push the egg through the Sp-Ut valve and into the Ut (Moive S1) (Yamamoto et al. 2006; Castaneda et al. 2020). Two sphincter muscles, the Sp neck and the Sp-Ut valve, can therefore block the passage of oocytes and promote their accumulation in the absence of sperm. We therefore specifically restored *daf-18* either in the Sp neck, or the Sp-Ut valve, of *fog-1; daf-18(ø)* doubles, using the *ipp-5* and *rsef-1* promoters, respectively (Bui and Sternberg 2002; Ghosh and Sternberg 2014). *Sp neck::DAF-18* rescued both oocyte arrest and GSC downregulation, while *Sp-Ut valve::DAF-18* did not (Figures 3B-3E). We conclude that DAF-18 activity in the Sp necks, each formed by 8 myoepithelial cells, is sufficient to non-autonomously promote oocyte arrest/accumulation in the absence of sperm, and to induce the concomitant downregulation in GSC proliferation.

### DAF-18’s lipid phosphatase activity is required to slow down germ tissue turnover

Since we previously found that DAF-16/FOXO, the forkhead transcription factor that mediates most long-term IIS effects (Ogg et al. 1997), was not required for oocyte accumulation and GSC downregulation in the absence of sperm (Narbonne et al. 2015; Qin and Hubbard 2015), we geared up to dissect the molecular mechanism by which DAF-18/PTEN was acting in the Sp neck to slow down oocyte turnover. DAF-18 is a dual-specificity phosphatase that could block ovulation through its lipid or protein phosphatase activity, or through a catalytically independent function.

To identify the key function, we introduced a G174E (equivalent to human G129E) substitution in endogenous DAF-18 to specifically remove its lipid phosphatase activity (Myers et al. 1997; Nakdimon et al. 2012; Chen et al. 2022; Wittes and Greenwald 2022). The resulting *fog-1; daf-18(G174E)* mutants wasted their oocytes like *fog-1; daf-18(ø)* (Figures 4A and 4B) while their GSCs showed sustained proliferation (Figures 4C and 4D). We conclude that DAF-18’s lipid phosphatase activity is required to promote oocyte and GSC quiescence in the absence of sperm.

**Figure 4.**
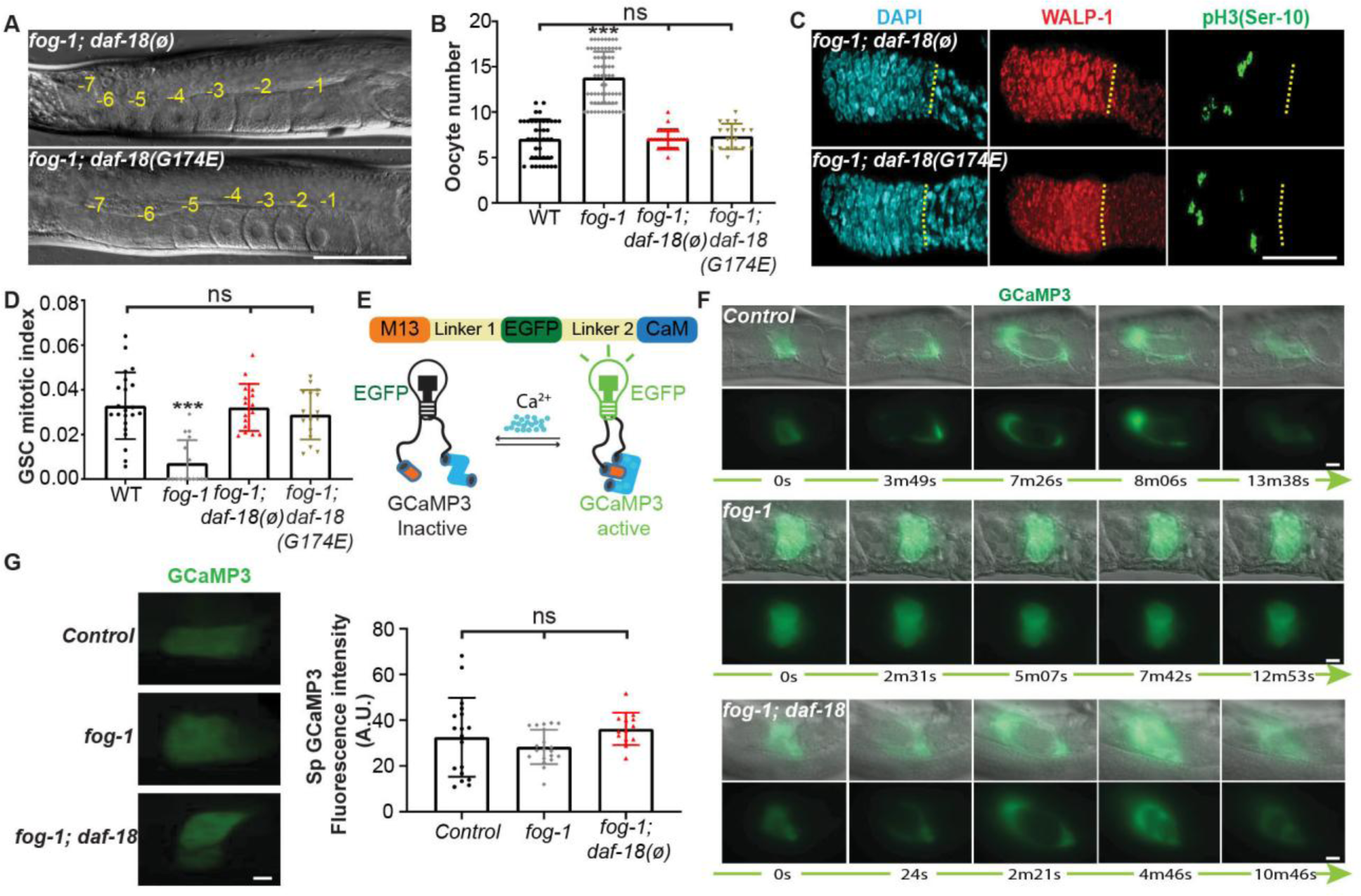
DAF-18’s lipid phosphatase activity slows down germ tissue turnover without affecting Ca^2+^ levels. **(A)** Representative DIC micrographs of A1 germlines of the indicated genotypes. Negative yellow numbers mark oocytes from distal to proximal. Dorsal, up; proximal, right ventral. **(B)** Average number (± standard deviation) of diakinesis-stage oocytes per gonad arm in A1 hermaphrodites of the indicated genotypes. Sample sizes: 42, 69, 30, 18. **(C)** Representative distal germlines dissected from A1 animals of the indicated genotypes, stained with DAPI (DNA; blue), anti-WALP-1 (proliferation maker; red) and anti-phospho[ser10] histone H3 (G_2_/M-phase marker; green). Yellow dotted lines, PZ boundary. **(D)** Average GSC MIs of A1 hermaphrodites of the indicated genotypes. Sample sizes: 22, 16, 16, 16. **(E)** Schematic representation of the GCaMP3 Ca^2+^ sensor, a synthetic fusion between EGFP, calmodulin (CaM) and the M13 peptide. When Ca^2+^ bound, the CaM domain interacts with the M13 alpha helix, resulting in bright fluorescence (Iseppon et al. 2022). **(F)** Representative time frames showing Sp Ca^2+^ flows during key ovulatory events in A1 hermaphrodites of the indicated genotypes (Supplementary Movies S1-S3). Sample sizes: 6, 13, 6. **(G)** Representative epifluorescence micrographs (left) and average Sp GCaMP3 fluorescence (right) of A1 hermaphrodites of the indicated genotypes. Sample sizes: 18, 20, 13. **(A, C, F, G)** Scale bars: 50 µm. **(B, D, G**) Asterisks indicate statistical significance *versus* all other samples. ns, not significant. **(C, F, G)** Distal, left.

### DAF-18 slows down germ tissue turnover by promoting Sp neck contractility, yet without influencing Ca^2+^ levels

DAF-18’s main lipid phosphatase activity dephosphorylates phosphatidylinositol (3,4,5)-triphosphate (PIP_3_) into phosphatidylinositol (4,5)-diphosphate (PIP_2_), which happens to be a substrate for the phospholipase C-ε PLC-1/PLC. PLC-1 cleaves PIP_2_ to generate inositol 1,4,5-triphosphate (IP_3_), which in turn opens the IP_3_-sensing ITR-1/IP_3_R Ca^2+^ channel that releases Ca^2+^ in the cytoplasm to trigger contractions (Kovacevic et al. 2013). We therefore hypothesized that DAF-18 could promote Sp neck contractility through increasing PIP_2_ levels and promoting PLC-1/ITR-1/Ca^2+^ signaling. To test this idea, we filmed ovulating hermaphrodites expressing the fluorescent Ca²⁺ sensor GCaMP3 in their spermatheca (Figure 4E) (Nakai et al. 2001; Bouffard et al. 2019). While Sp Ca^2+^ flows and ovulation occurred normally in sperm-bearing control hermaphrodites (Figure 4F; Movie S1), the Sp neck always remained constricted in spermless *fog-1* hermaphrodites and effectively prevented oocytes to enter the Sp (Figure 4F; Movie S2). In *fog-1; daf-18(ø)* mutants however, the Sp neck relaxed normally to allow oocyte entry as in sperm-bearing controls, after which the neck and bag contracted to expulse it into the Ut even though it was not fertilized (Figure 4F; Movie S3). Interestingly, we noticed that oocyte Sp entry time was significantly shorter in *fog-1; daf-18(ø)* mutants, revealing a lower Sp neck stretching resistance (Figure S3A; Movies S1-S3). We conclude that DAF-18 cell autonomously increases Sp neck contractility in the absence of sperm to restrict ovulation and arrest oocytes.

Contractility depends on actin/myosin interactions which are stimulated by rises in cytoplasmic Ca^2+^ concentrations (Wakabayashi 2015). We thus asked whether DAF-18 prevents Sp neck dilation by raising cytoplasmic Ca^2+^ levels. Sp GCaMP3 fluorescence intensities were however undistinguishable in control, *fog-1* and *fog-1; daf-18(ø)* hermaphrodites (Figure 4G), suggesting that DAF-18 promotes Sp neck contractility independently of Ca^2+^ levels. Yet, since the loss of *daf-18* is expected to reduce PIP_2_, which in turn forecasts reduced PLC-1/ITR-1/Ca^2+^ stimulation (Kariya et al. 2004), to firmly exclude this possibility, we still investigated the genetic interaction between *daf-18* and *plc-1*.

In the presence of sperm, the loss of *plc-1* causes the trapping of embryos in the Sp since its neck and bag lack the ability to strongly contract and expulse eggs through the Sp-Ut valve (Castaneda et al. 2020). Accordingly, the *fog-1; plc-1(ø)* double mutant exhibited oocyte trapping (Figure S3B). We reasoned that if DAF-18 was to affect Sp neck contractility exclusively through PLC-1, its removal should not exacerbate the *plc-1(ø)* phenotype. However, *fog-1; daf-18(ø); plc-1(ø)* triple mutants clearly had more Sp-trapped oocytes than *fog-1; plc-1(ø)* doubles (Figure S3B). Sp GCaMP3 fluorescence intensity also remained unfazed by these genetic perturbations (Figure S3C), confirming that the loss of DAF-18 does not affect Sp Ca^2+^ levels, even in the absence of PLC-1. Most strikingly, the loss of *daf-18* still largely prevented anovulated oocyte accumulation in the absence of *plc-1* (Figures S3B and S3D), suggesting that *daf-18* promotes their accumulation mostly independent of *plc-1*. We attribute the weak rescue of oocyte accumulation to the introduction of a new blockade in the path of oocytes in the absence of *plc-1*, this time at the Sp-Ut valve, that may have caused one or two oocytes to backlog into the proximal gonad once the Sp got maximally stretched (Figures S3B and S3D). Interestingly, the loss of *plc-1* had a negative effect on GSC proliferation on its own, an effect that was not additive with the negative homeostatic pressure present in *fog-1* mutants (Figure S3E). Germline feminization and the loss of *plc-1* may therefore negatively regulate GSC proliferation through the same downstream mechanism. Moreover, *fog-1; daf-18(ø); plc-1(ø)* triple mutants had a GSC MI that was intermediate between the *fog-1; plc-1(ø)* and the *fog-1; daf-18(ø)* doubles (Figure S3E), indicating that *daf-18* suppresses GSC proliferation in feminized hermaphrodites through a mechanism that is independent of *plc-1*.

To further explore the interaction between *daf-18* and Ca^2+^ signaling, we reduced the activity of the *C. elegans* unique IP_3_ receptor, ITR-1 (Baylis et al. 1999), using the viable *itr-1(sa73)* reduction-of-function *(rf)* allele (Dal Santo et al. 1999). We first note that feminized *itr-1(rf)* mutants had lower Sp intracellular Ca^2+^ levels than feminized *plc-1(ø)* mutants, but that this was again, unaffected by the loss of *daf-18* (Figure S3C). The stronger Ca^2+^ defect of *itr-1(rf)* versus *plc-1(ø)* may be explained by redundancy or compensation between *plc-1* and *plc-3* as the latter is also expressed in the spermatheca (Yin et al. 2004). Otherwise, the *itr-1(rf)* allele interacted with *daf-18(ø)* like *plc-1(ø)* in all assays (Figures S3B-S3G) except for one notable difference. Namely, and potentially due to their presumably looser Sp neck (Figures S3B and S3C), *fog-1; itr-1(rf)* doubles did not accumulate as many anovulated oocytes as *fog-1* singles (Figures S3B and S3F). As a result, the removal of *daf-18* in the *fog-1; itr-1(rf)* background had no additional effects on oocyte accumulation. But as seen with *plc-1(ø)*, *daf-18* still promoted GSC quiescence independently of *itr-1* (Figure S3G). Altogether, these data indicate that DAF-18 does not influence Sp Ca^2+^ levels and largely slow down oocyte turnover independently of PLC-1/ITR-1/Ca^2+^ signals. Oocyte accumulation downstream of *daf-18* nonetheless largely depended on the presence of *itr-1* (Figures S3B-S3G), the loss of which markedly reduced Sp intracellular Ca^2+^ levels (Figure S3C). Oocyte arrest and accumulation may therefore require ITR-1 activity for intracellular Ca^2+^ levels to reach a threshold that allows *daf-18* to strengthen the Sp neck.

### DAF-18 slows down germ turnover by reducing Sp PIP_3_ levels and AKT-1,2 activity

We reasoned that if Ca^2+^ levels were largely unfazed downstream of DAF-18’s product, the mutant’s ovulation defects may instead result from changes in substrate concentration. We thus first used anti-PIP_3_ antibodies to evaluate Sp levels. Quite strikingly, PIP_3_ was abundantly associated with Sp cell membranes, while the rest of the germline/gonad contained very little PIP_3_ (Figure 5A). Feminization did not perturb Sp PIP_3_ levels, but there were more PIP_3_ in the *daf-18(ø)* Sp, both in control and feminized hermaphrodites (Figure 5B). As DAF-18 activity lowers Sp PIP_3_ levels, we reasoned that it must in turn suppress AKT-1,2 (the *C. elegans* AKT/PKB orthologs) through reducing PDK-1/PDK1 activation (Murphy and Hu 2013).

**Figure 5.**
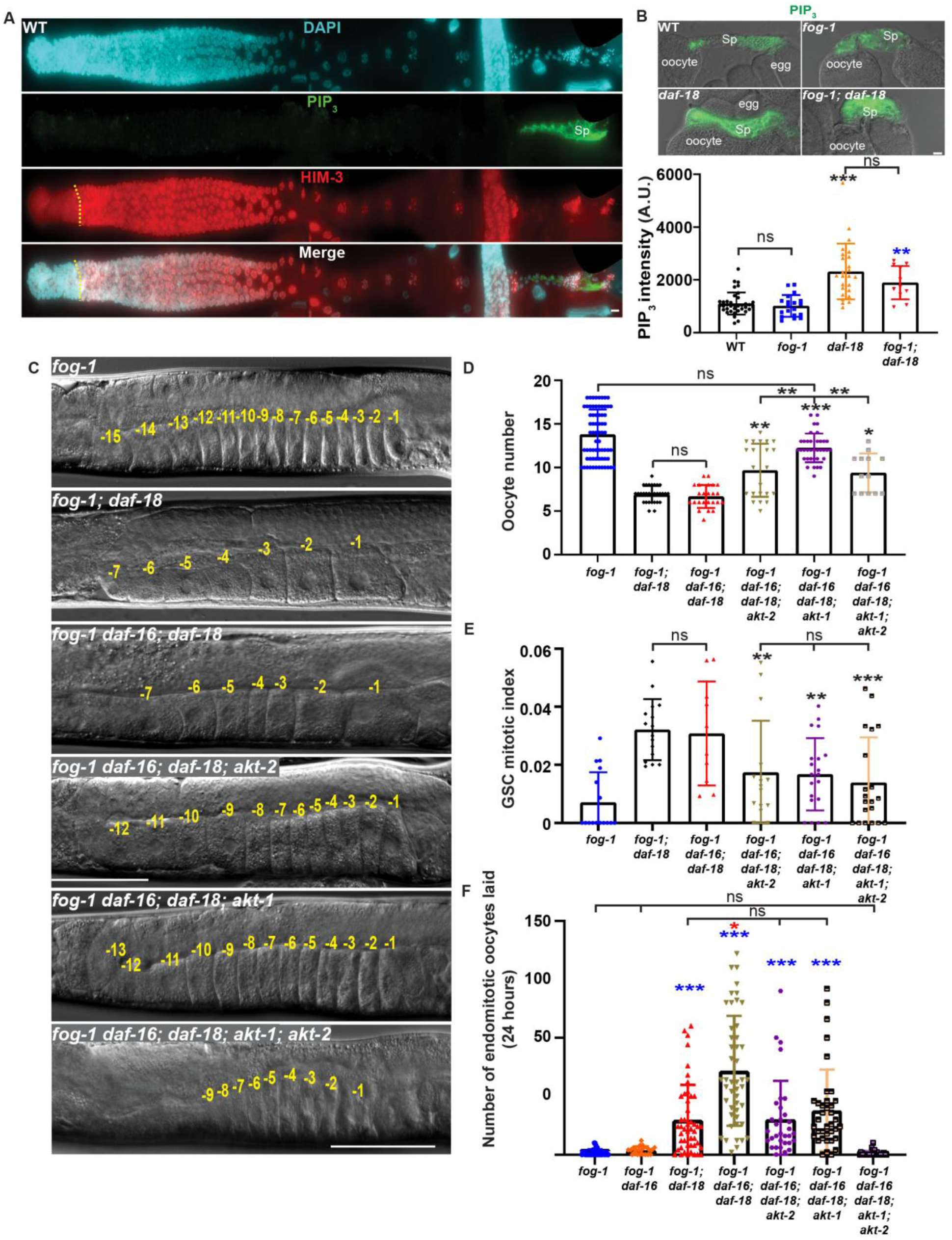
DAF-18 slows down germ turnover by reducing PIP_3_ levels and AKT-1,2 activity. **(A)** Representative epifluorescence micrographs of a whole germline dissected from a wild-type A1 hermaphrodite, stained with DAPI (DNA; blue), anti-PIP_3_ (PIP_3_; green), and anti-HIM-3 (differentiation marker, red). PIP_3_ is highly enriched at Sp cell membranes. Distal, left. **(B)** Representative DIC/epifluorescence micrographs of dissected proximal gonads of A1 hermaphrodites of the indicated genotypes, stained with anti-PIP_3_ (PIP_3_; green) (top) and average Sp PIP_3_ levels (bottom). Sample sizes: 38, 25, 19, 11. **(C)** Representative DIC micrographs of A1 germlines of the indicated genotypes. The DIC micrographs of *fog-1* and *fog-1; daf-18(ø)* controls are duplicates of Fig. 1C. Negative yellow numbers mark oocytes from distal to proximal. Dorsal, up; proximal, right ventral. **(D)** Average number (± standard deviation) of diakinesis-stage oocytes per gonad arm in A1 hermaphrodites of the indicated genotypes. Sample sizes: 68, 30, 24, 22, 36,13. **(E)** Average GSC MIs (± standard deviation) of A1 hermaphrodites of the indicated genotypes. Sample sizes: 16, 16, 10, 15, 20, 20. **(F)** Average number (± standard deviation) of endomitotic oocytes laid by hermaphrodites of the indicated genotypes over a period of 24 hours after A1. Sample sizes: 40, 22, 47, 47, 32, 37, 34. **(A-C)** Scale bars: 50 µm. **(B**,**D**-**F)** Asterisks indicate statistical significance versus the corresponding color-coded sample, or as indicated by brackets. The red asterisk represents statistical significance versus all other groups. ns, not significant.

Incidentally, *akt-1* and *akt-2* are highly expressed in the Sp (Padmanabhan et al. 2009), consistent with a prominent role in this tissue. However, *akt-1(ø)* and *akt-2(ø)* adults each had normal oocyte numbers and GSC MIs (Figures S4A-S4C). To investigate a potential redundancy, *akt-1/2(ø)* doubles were combined with a null mutation in the FOXO ortholog *daf-16* to prevent constitutive dauer formation and allow triples to grow into adults (Ogg et al. 1997). Both *fog-1 daf-16(ø)* doubles and *fog-1 daf-16(ø); akt-1/2(ø)* quadruples however had oocyte numbers and GSC MIs that matched those of *fog-1* singles (Figures S4A-S4C). Thus, in the absence of *daf-16*, *akt-1/2* do not significantly influence ovulation and GSC proliferation in normal or feminized hermaphrodites when *daf-18* is present. This is consistent with previous results that established *daf-18* as promoting oocyte arrest and GSC quiescence in spermless hermaphrodites independently of *daf-16* (Narbonne et al. 2015; Qin and Hubbard 2015).

We next investigated the loss of *akt-1/2* in the absence of *daf-18*. The removal of *daf-16* from *fog-1; daf-18(ø)* did not perturb oocyte numbers, nor the GSC MI (Figures 5C-5E). We then removed *akt-1* and/or *akt-2* from this triple mutant strain to generate *fog-1 daf-16(ø); daf-18(ø); akt-1(ø)* and *fog-1 daf-16(ø); daf-18(ø); akt-2(ø)* quadruples, as well as the *fog-1 daf-16(ø); daf-18(ø); akt-1/2(ø)* quintuple. Both quadruples had significantly more oocytes and lower GSC MIs than *fog-1; daf-18(ø),* and *fog-1 daf-16(ø); daf-18(ø)* controls (Figures 5C-5E), indicating that the loss of either *akt-1* or *akt-2* significantly rescues both *daf-18(ø)* defects. Conversely, an *akt-1* gain-of-function reduced oocyte accumulation and suppressed GSC downregulation in feminized hermaphrodites, partially phenocopying *daf-18(ø)* (Figures S4A-S4C). Although oocytes numbers in *fog-1 daf-16(ø); daf-18(ø); akt-1(ø)* quadruples were undistinguishable from those of *fog-1* controls (Figures 5D and 5E), these quadruples laid more endomitotic oocytes (Figure 5F), indicating that some unwanted ovulation still occurred.

Because of the partial suppression observed in each quadruple mutant and anticipated *akt-1/2* redundancy, we expected the simultaneous removal of *akt-1/2* to fully rescue feminized *daf-18(ø)* hermaphrodites. However, oocyte accumulation was poorly restored in the *fog-1 daf-16(ø); daf-18(ø); akt-1/2(ø)* quintuple compared with the *fog-1 daf-16(ø); daf-18(ø); akt-1(ø)* quadruple (Figures 5C and 5D). Yet, and for an unknown reason, the *fog-1 daf-16(ø); daf-18(ø); akt-1/2(ø)* quintuple mutant was somewhat sickly and grew slowly (Figure S4D), which may be linked to this partial rescue. Importantly however, the quintuple mutant stopped to futilely lay endomitotic oocytes (Figure 5F). These results together suggest that *akt-1/2* work additively to permit ovulation in the absence of sperm and *daf-18*, and that they are together sufficient to mediate the unwanted ovulations. The loss of DAF-18 therefore elevates PIP_3_ levels in the Sp neck and activate AKT-1,2 which, in turn and independently from DAF-16, promote Sp neck dilation to allow ovulation in the absence of sperm. The question therefore became how AKT-1,2 were suppressing Sp neck contractility to promote germ turnover, yet without affecting Sp Ca^2+^ levels or requiring DAF-16/FOXO.

### AKT-1,2 phospho-inhibit RHO-1/RHOA to inappropriately promote germ tissue turnover

Contractions happen when rises in intracellular Ca^2+^ activate a Ca^2+^-dependent myosin light chain (MLC) kinase that phosphorylates MLC and promotes its interaction with actin (Kelley et al. 2018). Yet, contractility is further modulated by the Rho-associated coiled-coil kinase LET-502/ROCK, which directly phosphorylates both MLC and MLC phosphatase to increase Ca^2+^ sensitivity (Kovacevic et al. 2013; Tan and Zaidel-Bar 2015). Hence, we reasoned that AKT-1,2 could affect Sp contractility through LET-502 without altering Ca^2+^ levels. We further deduced that AKT-1,2 were likely to act on LET-502’s upstream activator, the Ras homolog family member A (RHOA) ortholog RHO-1, or on LET-502 itself, since upstream players also influence Ca^2+^ levels (Castaneda et al. 2020).

We noticed a single perfectly conserved AKT RXRXXS/T consensus phosphorylation motif (Paradis and Ruvkun 1998) around RHO-1’s serine 73 (S73), but no such motif in LET-502 (Figure 6A). We used the High Ambiguity Driven protein-protein DOCKing (HADDOCK) (de Vries et al. 2010) platform to model a potential physical interaction between AKT-1’s kinase domain and RHO-1’s S73, and obtained a docking score predictive of its likelihood (Figures 6B-6D; Movie S4). To have a useful comparison, we repeated this *in silico* experiment between AKT-1 and DAF-16 to obtain the docking scores for each of its four well-established AKT target sites (Paradis and Ruvkun 1998). The AKT-1 docking score for RHO-1’s S73 (-112 ± 8.8) was highly negative suggesting that such interaction was favorable, with a value comprised within the range of those obtained for DAF-16’s sites (-80.7 to -139.7) (Figure 6B). Template-free docking scores between AKT-1 and RHO-1 or DAF-16 were also similar (Figure S5A) (Yan et al. 2020). To test whether this molecular interaction was possible, we performed an *in vitro* kinase assay between active human AKT-1 and human RHOA. The results confirmed that AKT-1 can directly phosphorylate RHOA (Fig. 6E).

**Figure 6.**
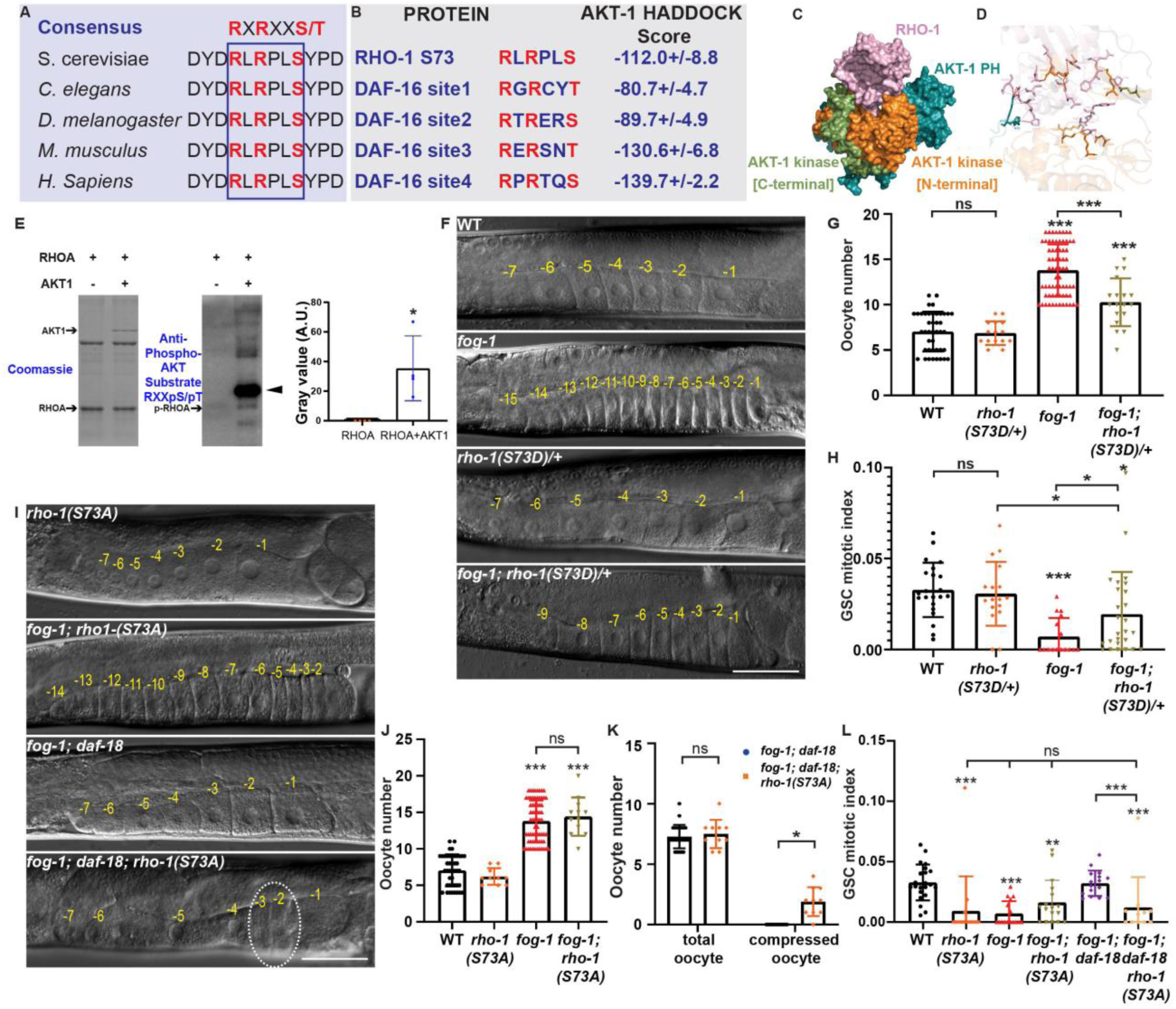
AKT-1,2 directly phospho-inhibit RHO-1/RHOA to promote germ tissue turnover. **(A)** A consensus AKT RXRXXS/T phosphorylation site lies within a perfectly conserved RHO-1/RHOA region. **(B)** The HADDOCK platform predicts a favorable docking energy between AKT-1’s kinase domain and the RHO-1(S73) interface, as for AKT-1 and its established DAF-16/FOXO target sites (Paradis and Ruvkun 1998). **(C)** HADDOCK generated AKT-1 (blue: PH domain; orange: kinase N-terminal; green: kinase C-terminal) RHO-1 (pink) complex. **(D)** Close-up of the AKT-1/RHO-1 interface showing predicted hydrogen bonds. **(E)** An *in vitro* kinase assay was performed between recombinant human RHOA and active recombinant AKT1 and Western blot analyses using an anti-phospho-Akt substrate antibody revealed that AKT1 can directly phosphorylate RHOA. **(F)** Representative DIC micrographs of A1 hermaphrodite germlines of the indicated genotypes. **(G)** Average number (±standard deviation) of diakinesis-stage oocytes per gonad arm in A1 hermaphrodites of the indicated genotypes. Sample sizes: 42, 15, 68, 18. **(H)** Average GSC MIs (±standard deviation) of A1 hermaphrodites of the indicated genotypes. Sample sizes: 22, 17, 16, 26. **(I)** Representative DIC micrographs of A1 germlines of the indicated genotypes. White dotted circle, visibly compacted oocytes. **(F, I)** Negative yellow numbers of mark oocytes from distal to proximal. Dorsal, up; proximal, right ventral. Scale bar: 50 µm. DIC micrographs of WT, *fog-1* and *fog-1; daf-18(ø)* controls are duplicates of Figure 1C. **(J)** Average number (±standard deviation) of diakinesis-stage oocytes per gonad arm in A1 hermaphrodites of the indicated genotypes. Sample sizes: 42, 10, 68, 12. **(K)** Although the *fog-1; daf-18(ø) rho-1(S73A)* strain does not accumulate oocytes, most animals have some visibly compacted oocytes, something that never occurs in *fog-1; daf-18(ø)* doubles. Sample sizes, 25, 10. **(L)** Average GSC MIs (±standard deviation) of A1 hermaphrodites of the indicated genotypes. Sample sizes: 22, 15, 16, 17, 16, 11. **(G-H, J-L)** Asterisks indicate statistical significance versus the corresponding color-coded sample, or as indicated by brackets. ns, not significant.

To evaluate whether AKT-1,2 may directly phosphorylate RHO-1 *in vivo* to promote Sp neck dilation, we mutated endogenous RHO-1’s S73 to either alanine (A) or aspartic acid (D), to respectively prevent or mimic phosphorylation (Leger et al. 1997; Bauer et al. 2003; Peng et al. 2012). Based on the *daf-18(ø)* phenotype and its suppression by *akt-1/2(ø)*, we predicted that a phospho-defective RHO-1(S73A) variant would no longer be inhibited by AKT and exhibit a persistent activity, while a phospho-mimetic RHO-1(S73D) variant would be constitutivel inhibited. Consistent with this, it was shown that such phosphorylation impaired the activity of Rac1, another member of the Rho family of small GTPases (Kwon et al. 2000).

Unfortunately, we could not obtain homozygous phospho-mimetic *rho-1(S73D)* animals as this substitution was embryonic lethal (Figures S5B and S5C), like *rho-1(ø)* (Fotopoulos et al. 2013). This result nonetheless suggests that constitutive phosphorylation at S73 would severely impair RHO-1 activity. We thus examined *fog-1; rho-1(S73D)/+* viable heterozygotes as their presumably halved functional RHO-1 levels may be insufficient to fully support Sp neck contraction and oocyte accumulation. Accordingly, *fog-1; rho-1(S73D)/+* heterozygotes accumulated fewer oocytes than *fog-1* controls and had an increased GSC MI (Figures 6F-6H). We conclude that in the absence of sperm and DAF-18, the presumed phosphorylation of RHO-1 by AKT-1,2 reduces RHO-1 activity and Ca^2+^ sensitivity to suppress Sp neck contractility, preventing any robust oocyte retention/accumulation and GSC downregulation.

Although they laid much fewer and poorly viable eggs (Figures S5D and S5E), homozygous *rho-1(S73A)* phospho-defective mutants were viable (Figure 6I). We expected that RHO-1(S73A) would prevent any AKT-1,2 inhibitory phosphorylation, and thus present a persistent activity, but not necessarily an increased or constitutive activity. Accordingly, the *rho-1(S73A)* mutation did not perturb oocyte numbers in the presence or absence of sperm (Figures 6I and 6J). We expected *rho-1(S73A)* to suppress ovulation caused by the loss of *daf-18* by preventing AKT-1,2-dependent RHO-1 downregulation. However, *fog-1; daf-18(ø) rho-1(S73A)* triple mutants did not accumulate more oocytes than *fog-1; daf-18(ø)* controls (Figures 6I and 6J). They nonetheless often exhibited visibly compressed oocytes (Figures 6I and 6K), indicating that their Sp neck opposed a greater resistance. This incomplete rescue may be linked to the substantial sterility of *rho-1(S73A)* mutants (Figures S5D and S5E), which could be unable to generate enough oocytes to sustain their accumulation. Consistent with this interpretation, *rho-1(S73A)* had a negative effect on GSC proliferation, suggesting that persistent RHO-1 activity limits GSC proliferation on its own (Figure 6L). Interestingly, feminization of *rho-1(S73A)* mutants did not further reduce GSC proliferation (Figures 6L), suggesting that feminization negatively regulates GSC proliferation by preventing RHO-1’s S73 phospho-inactivation. Moreover, GSC proliferation remained low when *daf-18* was removed from *fog-1; rho-1(S73A)* doubles, indicating that phosphorylation on RHO-1(S73) is required for AKT-1,2 to promote GSC proliferation in *fog-1; daf-18(ø)* mutants. Altogether, our data demonstrate that in the absence of sperm, DAF-18 activity suppresses AKT-1,2 and critically prevents them from directly phospho-inhibiting RHO-1. As a result, RHO-1 can enhance Sp neck Ca^2+^/contractility responses to suppress spontaneous ovulation. This allows unfertilized oocytes to accumulate in the proximal gonad and the homeostatic downregulation of germline stem cells to globally slow down germ tissue turnover (Figure 7A).

**Figure 7.**
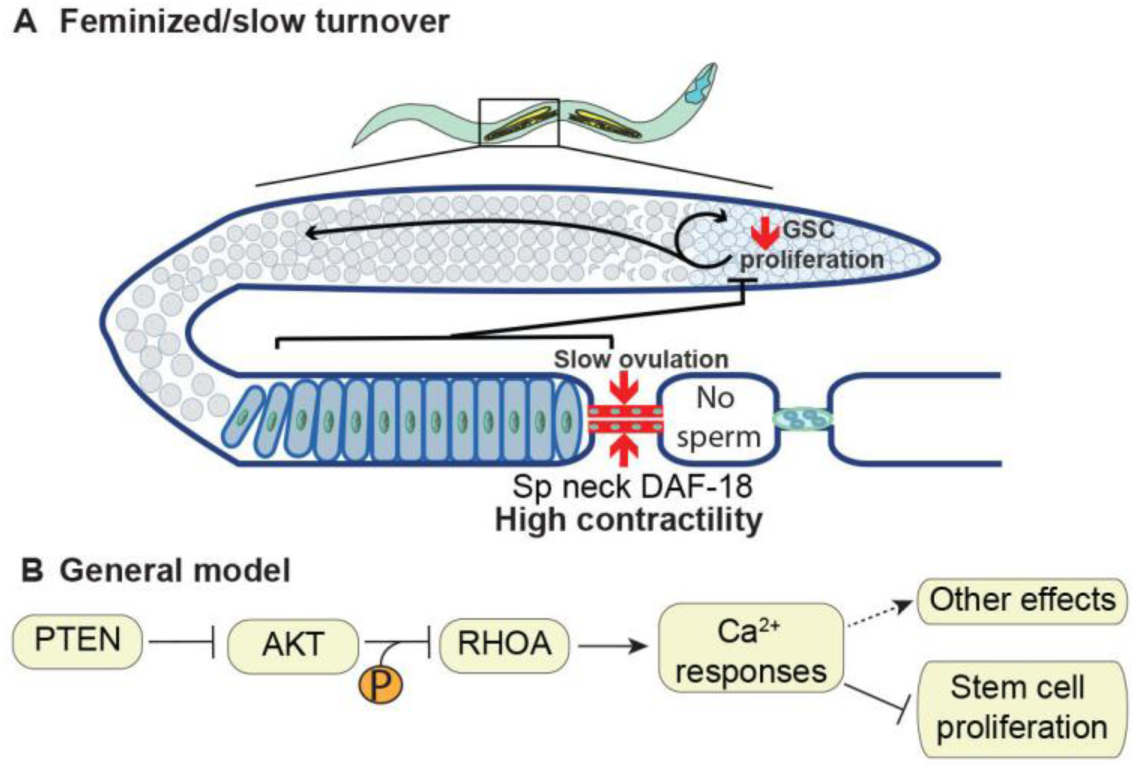
Models for how DAF-18/PTEN couples stem cell proliferation with differentiated cell needs. **(A)** In *C. elegans* hermaphrodites that lack sperm, DAF-18/PTEN critically suppresses AKT-1,2/AKT, thereby relieving its phospho-inhibition of RHO-1/RHOA. Activated RHO-1/RHOA thereby enhances Sp neck Ca^2+^ sensitivity, increasing its contractility to prevent spontaneous ovulation. This provokes the accumulation of unfertilized oocytes and the homeostatic feedback that downregulates GSC proliferation to globally slow down germ tissue turnover. **(B)** We extrapolate a general model in which PTEN could similarly prevent AKT from phospho-inhibiting RHOA to increase Ca^2+^ responses in differentiated cells and reduce stem cell proliferation. In *C. elegans*, the enhanced Sp neck contractility is what prevents oocytes from secreting an EGF-like growth factor LIN-3 that would otherwise promote GSC proliferation (Clandinin 1998), but in principle, the inhibition of stem cell proliferation could be accomplished more directly, for example if RHOA was enhancing the Ca^2+^-induced secretion of growth inhibitors by differentiated cells. Finally, this IIS-Ca^2+^ crosstalk could serve diverse other purposes.

## Discussion

We first note that the loss of DAF-18/PTEN did not provoke a noticeable imbalance in *C. elegans* germline homeostasis when IIS was activated and sperm was present. As such, as long as tissue needs remain elevated, IIS may be able to globally skew germline homeostasis towards a rapid tissue turnover rate in adults (Michaelson et al. 2010; Lopez et al. 2013) without any DAF-18/PTEN requirements. Such global regulation of tissue turnover by IIS may be conserved in mammals, where for example, it is known to act directly on osteoblasts to drive bone formation (Fulzele et al. 2010), but at the same time, enhances the capacity of osteoclasts to resorb bone matrix (Ferron et al. 2010). It will be interesting to find out whether PTEN activity is similarly dispensable in this context.

When IIS was activated but tissue needs were down however, as in well-fed spermless *C. elegans* hermaphrodites, we found that DAF-18/PTEN activity was critical to increase Ca^2+^ sensitivity in the differentiated somatic cells forming the Sp neck to increase their dilation resistance. This prevented oocyte activation/ovulation and thereby their secretion of the EGF-like growth factor LIN-3 (Clandinin et al. 1998), which would have otherwise promoted GSC proliferation, likely through activating MPK-1/ERK in adjacent somatic gonadal sheath cells (Robinson-Thiewes et al. 2021). Upon removing DAF-18/PTEN from spermless hermaphrodites, germline homeostasis was therefore completely uncoupled as the Sp neck Ca^2+^ sensitivity remained low despite the lack of sperm/oocyte demand and as a result, oocytes were spontaneously ovulated and elevated GSC proliferation futilely maintained a full-scale oogenesis program, only to jettison all resulting oocytes. If such a homeostatic imbalance was to occur in a tissue that does not actively expulse its terminally-differentiated cells, like an epithelium for example, we predict that it would cause a hyperaccumulation of differentiated cells. And indeed, preventing oocyte laying by removing the redundant OMA-1,2 zinc finger proteins that permit oocyte maturation (Detwiler et al. 2001) in *daf-18(ø)* mutants provoked a disorganized hyperaccumulation of oocyte-like cells, which was caused by ongoing, homeostatically unchecked GSC proliferation (Figure. S6) (Narbonne et al. 2017; Valet and Narbonne 2022). Consistent with this, mice and humans born hemizygous for PTEN have a predisposition to develop similar disorganized hyperaccumulations of differentiated cells, called hamartomas (Ali and Mulita 2026), in multiple tissues (Liaw et al. 1997; Di Cristofano et al. 1998). Our work therefore suggest that such differentiated benign tumors may arise from a homeostatic uncoupling between stem/progenitor cell proliferation rates and differentiated cell needs. If the new principles that were discovered here in *C. elegans* are conserved, we further predict that in PTEN hamartoma tumor syndrome patients, periods during which differentiated cell needs are high may not be problematic, but that it may be specifically during periods with reduced differentiated cell needs that reduced PTEN activity provokes tumorigenesis.

This work’s main advance might however lie in the identification of the cell and molecular mechanisms that underlie the formation of differentiated hamartoma-like tumors. It was striking that the presence of DAF-18/PTEN within the stem cells at the base of the germ hamartomas did not matter at all. Indeed, and although there is overwhelming evidence that the loss of PTEN in stem cells confers a cell autonomous proliferative advantage (Groszer et al. 2006; Wang et al. 2006; Hill and Wu 2009; Magee et al. 2012), our results emphasize that the generation of hamartoma-like tumors may require tissue-level stem cell proliferation homeostasis deregulation.

Despite the peculiar architecture of the *C. elegans* germline, with its turnover rates relying on another tissue’s contractility status, we speculate that the loss of PTEN could skew differentiated cell turnover more directly in many other tissue types, through influencing growth factor secretion by differentiated cells. Indeed, since RHOA activity also modulates exocytosis and the secretion of diverse factors (van der Burgh et al. 2014; Ng et al. 2022; Wang et al. 2022), its activity could more directly influence the release of stem cell regulatory factors by differentiated cells in tissues where their turnover rates are either self-regulated or regulated by distant organs.

In addition to the perfect conservation of the AKT site on RHOA and its *in vitro* validation using the human proteins, there is *in vivo* evidence that PTEN activity promotes Ca^2+^ sensitivity in mammalian cells. Notably, a cardiomyocyte-specific PTEN deletion in mice caused a dramatic decrease in cardiac contractility that depended on PI3Kγ (Crackower et al. 2002). Moreover, type II diabetes is accompanied by an increase in the contractility of peripheral blood capillaries (Okon et al. 2005) that may result from a reduction in AKT-mediated RHOA inhibition due to insulin resistance. We therefore suspect that a negative regulation of contractility by AKT occurs in human tissues. We thus propose that PTEN ensures the homeostatic balance of adult tissue size and prevent hamartoma tumors across organisms by preventing AKT from phospho-inhibiting RHOA and hence, Ca^2+^-sensitizing key differentiated cells under reduced differentiated cell needs, while this direct phosphorylation may play a role in a variety of other biological contexts (Figure. 7B).

## Materials and Methods

### C. elegans genetics

Nematodes were maintained at 15℃ on standard NGM plates and fed *E. coli* bacteria of the strain OP50, unless otherwise indicated (Brenner 1974). The Bristol isolate (N2) was used as wild type throughout. All strains, alleles, deficiencies and transgenes used are listed in Table S1.

### Plasmids and transgenics

We used the Gibson method (Gibson et al. 2009) for assembling all plasmids. The source DNA and primers that were used, as well as microinjection concentrations, are listed in Table S2.

Extra-chromosomal arrays were generated by standard germline microinjections at a total concentration of 200 ng/μL, using pKSII as filler DNA and pCFJ104[*Pmyo-3::mCherry*] (5 ng/μL) or pMR352[*Pmyo-2::GFP*] (50 ng/μL) as co-injection markers (Mello et al. 1991; Frokjaer-Jensen et al. 2008; Narbonne and Roy 2009). For *Punc-54*, we used the fragment driving enhanced muscle expression previously coined “*PEunc-54*” (Masse et al. 2005). To rescue DAF-18 specifically in the germline, we used CRISPR/Cas9 to insert a single copy of a *Pmex-5::GFP::DAF-18(+)* + *unc-119(+)* fragment at ttTi5605 (LG II, +0.77 M.U.) into *unc-119(ed3)* mutants, using pPOM4 (Table S2) and pDD122 (Dickinson et al. 2013). A single line (*narSi5*) was obtained after injection of > 80 hermaphrodites. To generate the *daf-18(G174E), rho-1(S73A)* and *rho-1(S73D)* variants, we used the *dpy-10* co-CRISPR strategy with the Paix *et al*. protocol(Arribere et al. 2014). We further grew P0s on *cku-80(RNAi)* to favor repair by homologous recombination for all CRISPR/Cas9 genomic editions (Ward 2015). The sgRNAs and repair templates sequences are listed in Table S3.

### Oocyte counts

Late-L4 stage hermaphrodites were transferred from 15℃ to a new plate at 25℃ and after 3 days, F1 late-L4s hermaphrodites were synchronized by picking them to a new plate at 25℃ based on vulva development (Seydoux et al. 1993). They were grown for an additional 24 hours (Narbonne et al. 2015). This procedure allowed to inactivate the temperature sensitive *fog-1(q253)* allele throughout F1 larval development to prevent sperm formation (Barton and Kimble 1990). Resulting feminized day 1 adults (A1) were harvested, paralyzed with 0.1% Tetramisole (Sigma, L9756) and mounted onto M9 + 3% agarose pads. The number of oocytes per gonad arm, and their compaction status, were determined by differential interference contrast (DIC) examination.

### Germline mitotic index

Progenitor zone (PZ) mitotic indexes were evaluated as previously described (Crittenden et al. 2006; Narbonne et al. 2015; Narbonne et al. 2017; Hubbard and Schedl 2019; Robinson-Thiewes et al. 2021). Synchronized A1 hermaphrodites were generated as above, and their gonads were dissected and stained as previously described (Robinson-Thiewes et al. 2021). Briefly, hermaphrodites were transferred in an 8 µL drop of 1X PBS on a microscope slide cover glass and quickly dissected using a 25G surgical needle tip. The cover glass was then flipped onto a poly-L-lysine coated slide and submitted to a dry-ice freeze-crack procedure. Samples were fixed in -20℃ methanol for 1 minute and postfixed in a 3.7% paraformaldehyde solution (3.7% paraformaldehyde, 1X PBS, 0.08 M HEPES, 1,6 mM MgSO_4_ et 0,8 mM EGTA) for 30 minutes. Using 1ml PBST (PBS + 0.1% Tween 20) each time, slides were rinsed twice followed by a 10-minute incubation in PBST. Samples were then covered with 300 µL blocking solution (PBST + 3% BSA) for 1h at room temperature. Samples were stained using primary rabbit anti-WAPL-1 (1:500, Novus Biologicals #49300002) to mark GSCs along with their proliferating progeny (Kocsisova et al. 2018) and mouse anti-phospho[ser10]-histone H3 antibodies (1:250, Cell Signaling Technology #9706) to mark G2/M-phase nuclei, and counter-stained with 0.7 µg/mL 4’6-diamidino-2-phenylindole (DAPI) to highlight all nuclei. PZ nuclei counting in 3 dimensions was partially automated using an ImageJ plugin developed by Dr Jane Hubbard’s laboratory (Korta et al. 2012).

### Image acquisition and processing

For Figures 1C, 2A, 3B, 4A, 5D, 6E, 6H, S2A, S2B, S3B, S4A, DIC images and/or epifluorescence z-stacks were acquired every micron using a Plan-Apochromat 20x dry objective (NA 0.8) mounted on an inverted Zeiss Axio Observer.Z1. Micrographs were stitched using the Zen software (2.6 Blue Edition) and animals were straightened for ease of visualization using ImageJ. Epifluorescence signals were overlaid to the DIC micrographs using ImageJ.

For the high-resolution confocal fluorescence acquisitions in Figures 1A, 4G and S3C, A1 hermaphrodites were anaesthetized and mounted for imaging as above. Coverslips were sealed with VALAP (1:1:1 vaseline, lanolin, paraffin), and 0.35 μm z-step stacks were acquired using a Leica SP8 point scanning confocal microscope with an HC PL APO CS2 63x/1.30 numerical aperture oil immersion objective.

For Figure 4F and Movies S1-S3, z-stack DIC, epifluorescence micrographs and Movies were acquired every micron using a Plan-Apochromat 40x/1.4 oil objective mounted on an inverted Zeiss Axio Observer.Z1. Epifluorescence signals were overlaid to the DIC micrographs using ImageJ. All DIC micrographs show single focal planes, and where applicable, overlaid fluorescent micrographs also show the corresponding single focal plane.

Figure 5A micrographs were acquired every micron using a 60x oil immersion objective mounted on a DeltaVision microscope, maximally projected, stitched, straightened and thresholded using ImageJ.

For GSC MI evaluations (Figures 1D, 2C, 3D and 4C), representative maximal projections from z-stacks epifluorescence micrographs of the distal gonad acquired every micron using a Plan-Apochromat 40x/1.4 oil objective mounted on an inverted Zeiss Axio Observer.Z1 are shown.

### DAF-18::mNG quantification

Synchronized A1 hermaphrodites were generated, paralysed and mounted on a M9 + 3% agarose pad and imaged as above. Z-stacks were merged into single images using maximum intensity projection in ImageJ. A 10-pixel-wide segmented line was manually drawn along the membrane separating the −1 and −2 oocytes to obtain the mean fluorescence intensity, which was then background-subtracted.

### Dauer formation assays

Dauer formation was scored as described in previous reports (Paradis and Ruvkun 1998). Briefly, for Figure 1G, bleached batches of eggs were allowed to hatch in the absence of food at 15℃ for 36 hours and synchronized L1s were plated at 25°C to induce dauer entry. Dauer formation rate was calculated as the number of dauers divided by the total number of larvae after 96 hours. For Figures 2A and 2B however, hermaphrodites were picked together with males at the L4 stage and allowed to mate for 24 hours. Hermaphrodites were then singled onto new plates at 25°C and dauer formation rates in the crossed F1 progeny, or F2 progeny from crossed F1s, were evaluated.

### Ca^2+^ imaging and quantification

Intracellular Sp Ca^2+^ was evaluated in A1 hermaphrodites using the *Pfln-1::GCaMP3 in vivo* sensor (a kind gift from Erin J. Cram) (Kovacevic et al. 2013). Fluorescence intensity profiles were obtained by confocal microscopy and processed using ImageJ as previously described (Kovacevic et al. 2013). A segmented line was used to define a polygonal region of interest (ROI) corresponding to the Sp. Mean fluorescence intensity was measured for each region of interest (ROI), followed by background subtraction.

### PIP_3_ staining and quantification

A1 hermaphrodites were harvested, and their gonads were dissected and stained as described for GSC MI evaluations above (Robinson-Thiewes et al. 2021). Primary mouse monoclonal anti-PIP_3_ antibodies (1:100, Echelon Z-P345) and rabbit anti-HIM-3 (a kind gift from M. Zetka), and secondary A488-conjugated goat anti-mouse or A546-conjugated goat anti-rabbit antibodies (both at 1:500, Invitrogen Cat# A-32731 and A-11035, respectively) were used. DAPI was used as a counterstain. For quantification, micrographs were transferred to Imaris 9.2.1, and based on the green channel (A488), 3D models were individually thresholded to best fit the spermatheca contour. For PIP_3_ quantification, the GFP object was selected, and using the edit tool, its mean intensity was pulled from the statistics tab, and background subtracted.

### Endomitotic oocyte laying quantification

Synchronized A1 hermaphrodites were generated as above and singled to a new plate at 25°C for an additional 24 hours. Adults were removed and the number of endomitotic oocytes laid during this 24-hour period was determined under a dissecting microscope.

### Developmental milestone analysis

Developmental milestone analysis was scored as described in previous reports (Woodruff et al. 2019). Briefly, a large population of *C. elegans* was bleach-synchronized as described above. L1s were placed on seeded plates at 25°C and monitored every 24 hours. Developmental progression was tracked by counting the number of larvae that had reached, or developed beyond, the L4 stage. Individuals were plotted by their developmental status (“0” = yet to reach milestone; “1” = reached milestone).

### *In silico* protein modelling and docking predictions

The predicted ternary structures of AKT-1 and RHO-1 were generated using SWISS-MODEL (Guex et al. 2009; Waterhouse et al. 2018). Their amino acid sequences were aligned against the Protein Data Bank, and the crystal structures 1H10 and 3CQW were selected as templates to fold AKT-1 and RHO-1, respectively. We used the High Ambiguity Driven Protein-Protein DOCKing (HADDOCK) 2.4 (de Vries et al. 2010) platform to predict the physical interaction between the AKT-1 kinase domain and RHO-1’s S73 site, and to obtain docking scores predictive of the likelihood of such an interaction. The HDOCK platform was also used to confirm favorable docking scores.

### *In vitro* kinase assay and western blotting

*In vitro* kinase assays were performed essentially as described previously (Narbonne and Roy 2009). Briefly, 0.15 μg of active recombinant human AKT1 protein (0.1 μg/μL; Cat# A16-10G, SinoBiological) was incubated with 1 μg of recombinant human RhoA protein (His-tag; Cat# NBP1-50933, Novus) in Kinase Assay Buffer III (200 mM Tris-HCl pH 7.4, 100 mM MgCl₂, 0.5 mg/mL BSA; Cat# K03-09, SinoBiological). Fresh DTT was added into the Kinase Assay Buffer III, at a final concentration of 0.25 mM, and ATP (Cat# A50-09, SinoBiological) was added at a final concentration of 0.04 nM. Reactions lacking AKT1 were used as negative controls. All reactions were carried out at room temperature for 40 min and terminated by the addition of SDS loading buffer, followed by boiling at 95°C for 8 min.

Reaction products were analyzed by Western blots as described previously (Deng et al. 2019). Membranes were probed with phospho-Akt substrate (RXXpS/pT) (1:1000; clone 110B7E, Cell Signaling Technology) or RHOA antibody (1:1000; clone 2117, Cell Signaling Technology) and detected using HRP-conjugated goat anti-rabbit IgG (1:2000) (Cat#1705046 BIO-RAD) and ECL (Cat# 170-5061 BIO-RAD).

### Statistical analysis

Sample normality was verified using the Shapiro-Wilk test. Variance was verified with the F test for pairwise comparisons or with the Brown-Forsythe test for datasets comprising more than two groups. Tests used are indicated in the figure legends and were chosen according to the following criteria. For pairwise comparisons with normal distributions and equal variances, we used the unpaired two-tailed t-test. For non-gaussian distributions, the Mann-Whitney test was instead used. When the distribution was normal, but the variance was unequal, we used Welch’s t-test. For multi-group comparisons that all had normal distributions and equal variances, we used the one-way ANOVA with Tukey’s multiple comparisons. For non-gaussian distributions, we used the Kruskal-Wallis with Dunn’s multiple comparisons. When the distribution was normal, but the variances were unequal, we used the Brown-Forsythe and Welch’s ANOVA with Dunnett’s T3 multiple comparisons. Graphs were generated using GraphPad Prism 8. Asterisks indicate statistical significance (***: P<0.001; **: P<0.01; *: P<0.05) to all other samples unless otherwise specified. Statistical details, including all sample sizes, are indicated in the figure legends.

## Competing interests

Authors declare that they have no competing interests.

## Acknowledgments

We thank Jean-Claude Labbé, Alexandre Fisette and Erwan Pernet for constructive comments and edits on the manuscript; Erin J. Cram, Monique Zetka and Florence Solari for reagents and strains; the *Caenorhabiditis* Genetics Center (CGC) and WormBase for their essential roles in *C. elegans* research.

## Author Contribution

JD: Study design, experimentation, data analysis, presentation, manuscript drafting. ICDM: Helped generating UTR724, contributed to Figure S4B and S4C.

AMC: Helped generating UTR507, contributed Fig. 6B-6D and Movie S4.

OG: pOG1 design and construction, corresponding lines derivation and preliminary analyses; contributed to Figure 3B.

VR: Key preliminary analyses towards Figure 1G and 2A. POM: pPOM4 design and construction.

MJS: Supervision and manuscript editing.

PN: Funding, study design, experimentation, supervision, manuscript drafting and editing.

## Funding

Natural Sciences and Engineering Research Council of Canada (NSERC) (RGPIN-2019-06863, RGPAS-2019-00017, DGECR-2019-00326, RGPIN-2026-05498)

Canadian Institutes of Health Research (CIHR) (PJT-169138) Fondation Marcel et Rolande Gosselin

Fonds de recherche du Québec (FRQS) J2 bursary scholar (310643)

